# APEX2 proximity labeling of RNA in bacteria

**DOI:** 10.1101/2024.09.18.612050

**Authors:** Hadi Yassine, Elizabeta Sirotkin, Omer Goldberger, Vincent Lawal, Daniel B. Kearns, Orna Amster-Choder, Jared M. Schrader

**Affiliations:** Department of Biology, Indiana University, Bloomington, IN, 47405, USA; Departments of Chemistry and Biological Sciences, Wayne State University, Detroit, MI 48202, USA; Department of Microbiology and Molecular Genetics, IMRIC, The Hebrew University Faculty of Medicine, Jerusalem 91120, Israel

## Abstract

Rapid spatially controlled methods are needed to investigate RNA localization in bacterial cells. APEX2 proximity labeling was shown to be adaptable to rapid RNA labeling in eukaryotic cells, and through the fusion of APEX2 to different proteins targeted to different subcellular locations, has been useful to identify RNA localization in these cells. Therefore, we adapted APEX2 proximity labeling of RNA to bacterial cells by generating an APEX2 fusion to the RNase E gene, which is necessary and sufficient for BR-body formation. APEX2 fusion is minimally perturbative and RNA can be rapidly labeled on the sub-minute timescale with Alkyne-Phenol, outpacing the rapid speed of mRNA decay in bacteria. Alkyne-Phenol provides flexibility in the overall downstream application with copper catalyzed click-chemistry for downstream applications, such as fluorescent dye-azides or biotin-azides for purification. Altogether, APEX2 proximity labeling of RNA provides a useful method for studying RNA localization in bacteria.

**Motivation:** Studies over the past several years have shown that distinct RNAs can be targeted to subcellular locations in bacterial cells. The ability to investigate localized RNAs in bacteria is currently limited to imaging-based approaches or to laborious procedures to isolate ribonucleoprotein complexes by grad-seq, HITS-CLIP, or Rloc-seq. However, a major challenge in studying mRNA localization in bacterial cells is that bacterial mRNAs typically last for only a few minutes in the cell, while experiments to investigate their localization or interaction partners can take much longer. Therefore, rapid methods of studying RNA localization are needed to bridge this technical challenge.

**Highlights:**

- APEX2 proximity labeling can be applied to RNA in bacteria
- APEX2 RNA labeling reactions occur on the sub-minute timescale.
- APEX2 workflow requires less material and time than current methods.
- Alkyne-Phenol APEX2 substrate provides flexibility with click-chemistry.

## Introduction

Bacterial mRNAs have been found to localize to distinct subcellular locations. At a global level, many mRNAs have been found to be associated with the nucleoid, membrane, or cell poles^1–4^ and it has been hypothesized that mRNA localization can be important for proper gene expression. For example, Flagellin mRNA in Campylobacter jejuni localizes to the cell poles which may facilitate cotranslational flagellar assembly^5^. In addition, biomolecular condensates, which are non-membrane bound organelles often assembled through phase separation, have been rapidly expanding in bacterial cells, with multiple involved in RNA metabolism^4,6^. Yet the functional significance of localization of RNAs to biomolecular condensates in bacteria has just started to be explored.

GRAD-seq, HITS-CLIP, and Rloc-seq methods^1,7,8^ have been successful at identifying RNAs associated with specific RNPs or localized to certain subcellular locations. However, a major difficulty is that bacterial mRNAs are very short lived, and these methods require labeling or isolation times much longer than a typical mRNAs half-life, leading to the potential for false negatives. Therefore, rapid methods are needed to isolate the associated labile bacterial mRNA before they are degraded.

APEX2 proximity labeling allows for spatially controlled and rapid RNA labeling^9–12^, making it a useful method to assay for localized bacterial RNAs. In this manuscript, we show that APEX2 proximity labeling can be adapted to bacterial condensates via fusion of the core BR-body scaffold RNase E to APEX2. RNA can be labeled robustly by APEX2, as many bacteria natively produce heme, including E. coli, B. subtilis, and C. crescentus. With the addition of H_2_O_2_ and Alkyne-Phenol triggering labeling, allowing copper catalyzed click chemistry of Alkyne-Phenol labeled RNA with a variety of azides such as Cy5-azide. This protocol can be used to rapidly label bacterial RNAs by APEX2 on the sub-minute timescale, and which can be rapidly purified via copper catalyzed click chemistry conjugation of biotin-peg3-azide followed by streptavidin purification. Through the fusion of APEX2 to different localized RNA binding proteins in bacteria, APEX2 proximity labeling has the potential to improve our knowledge of RNA localization in bacteria.

## Materials and Methods

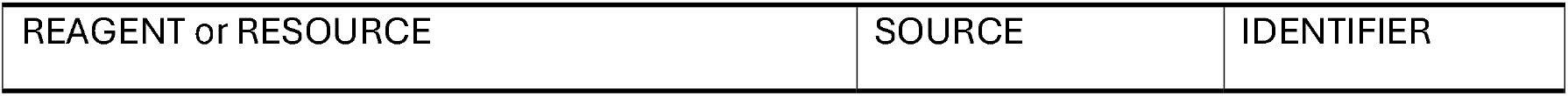
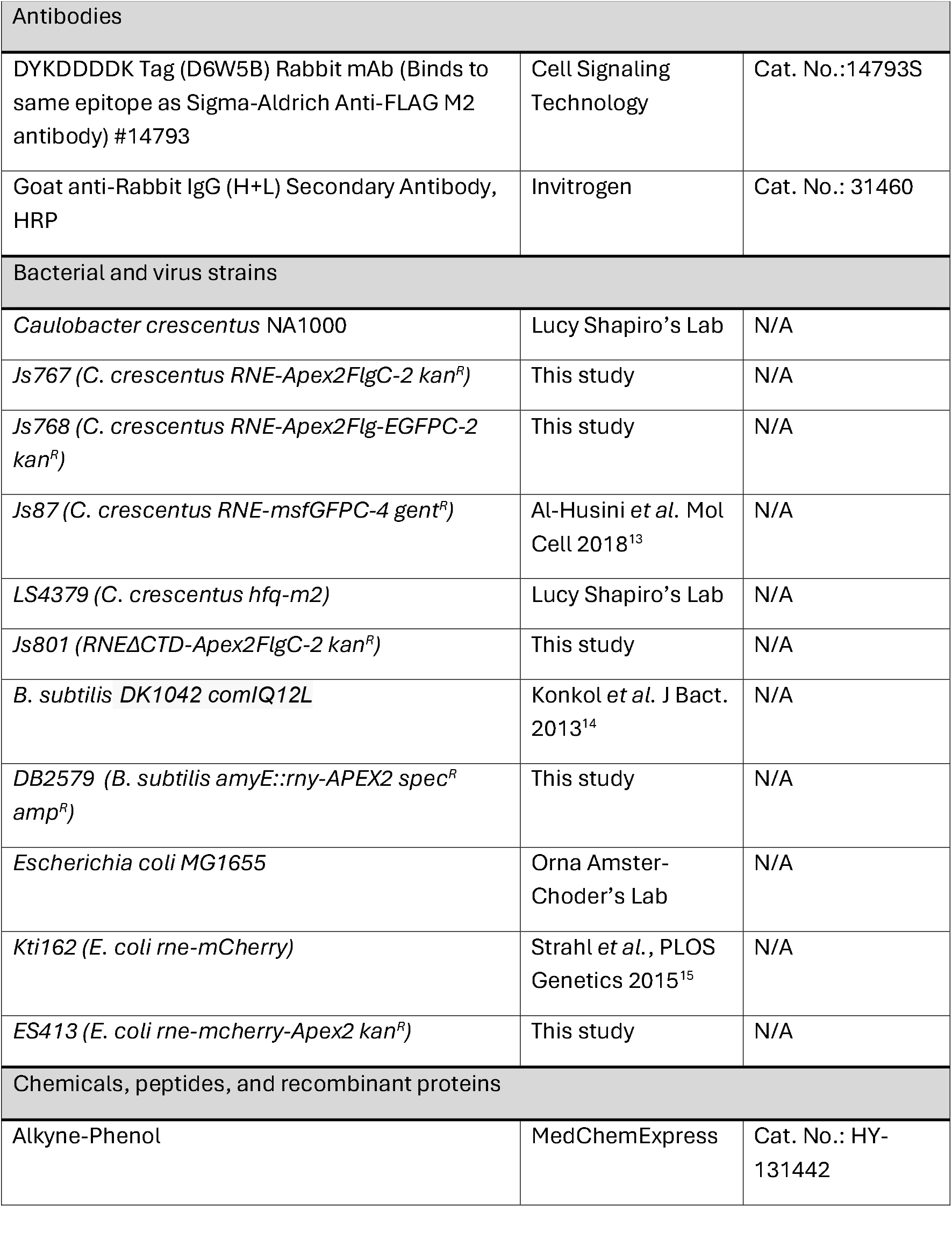
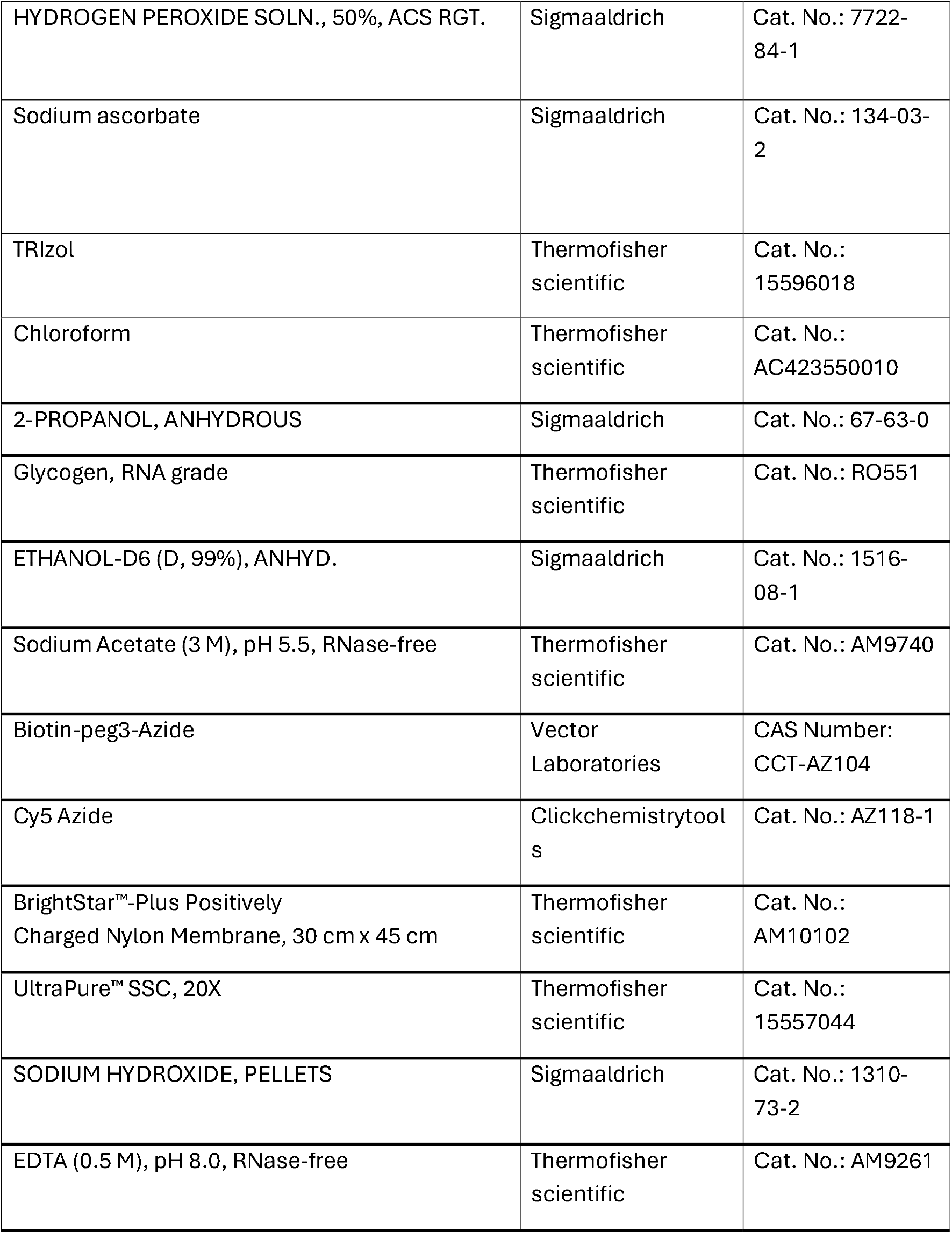
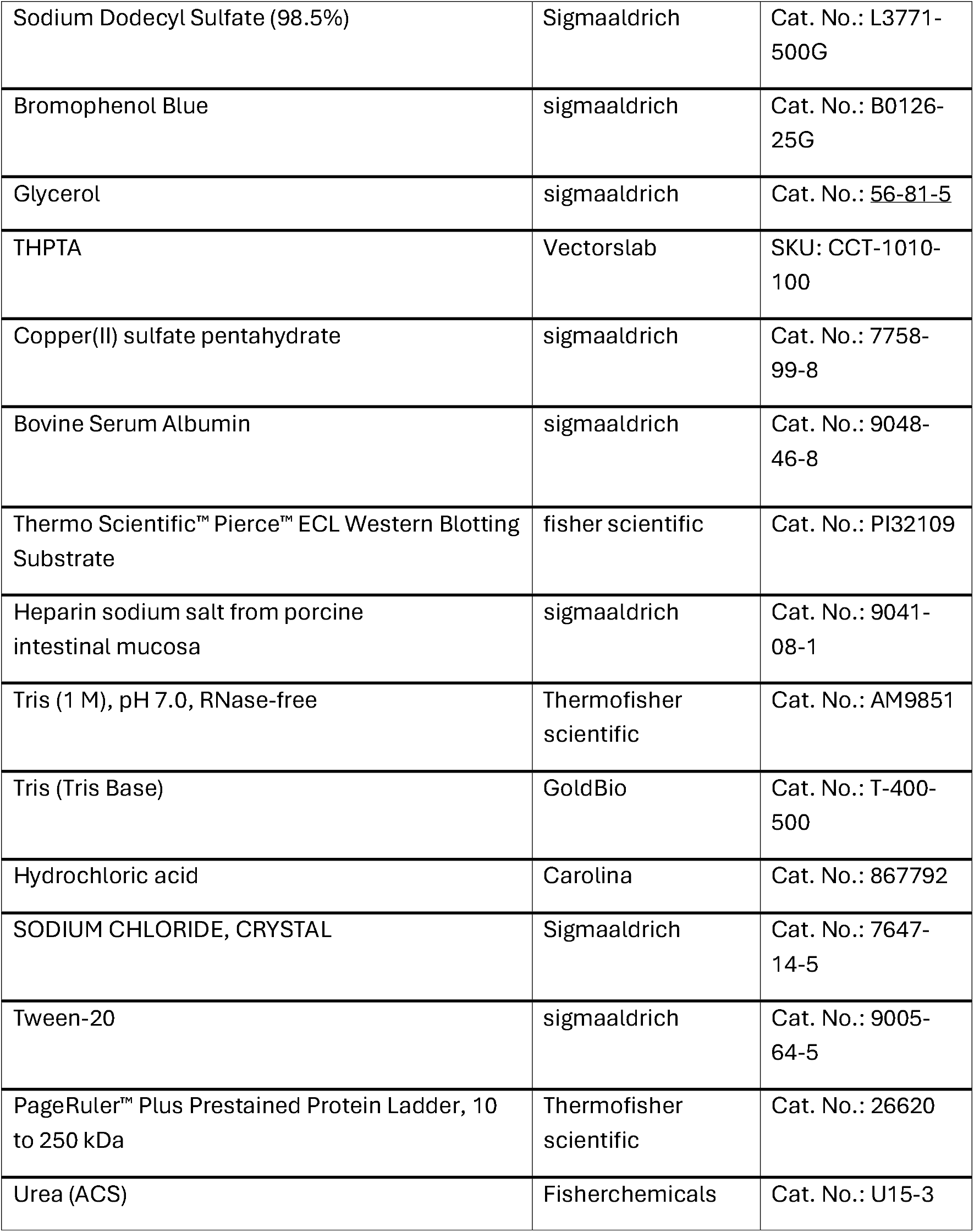
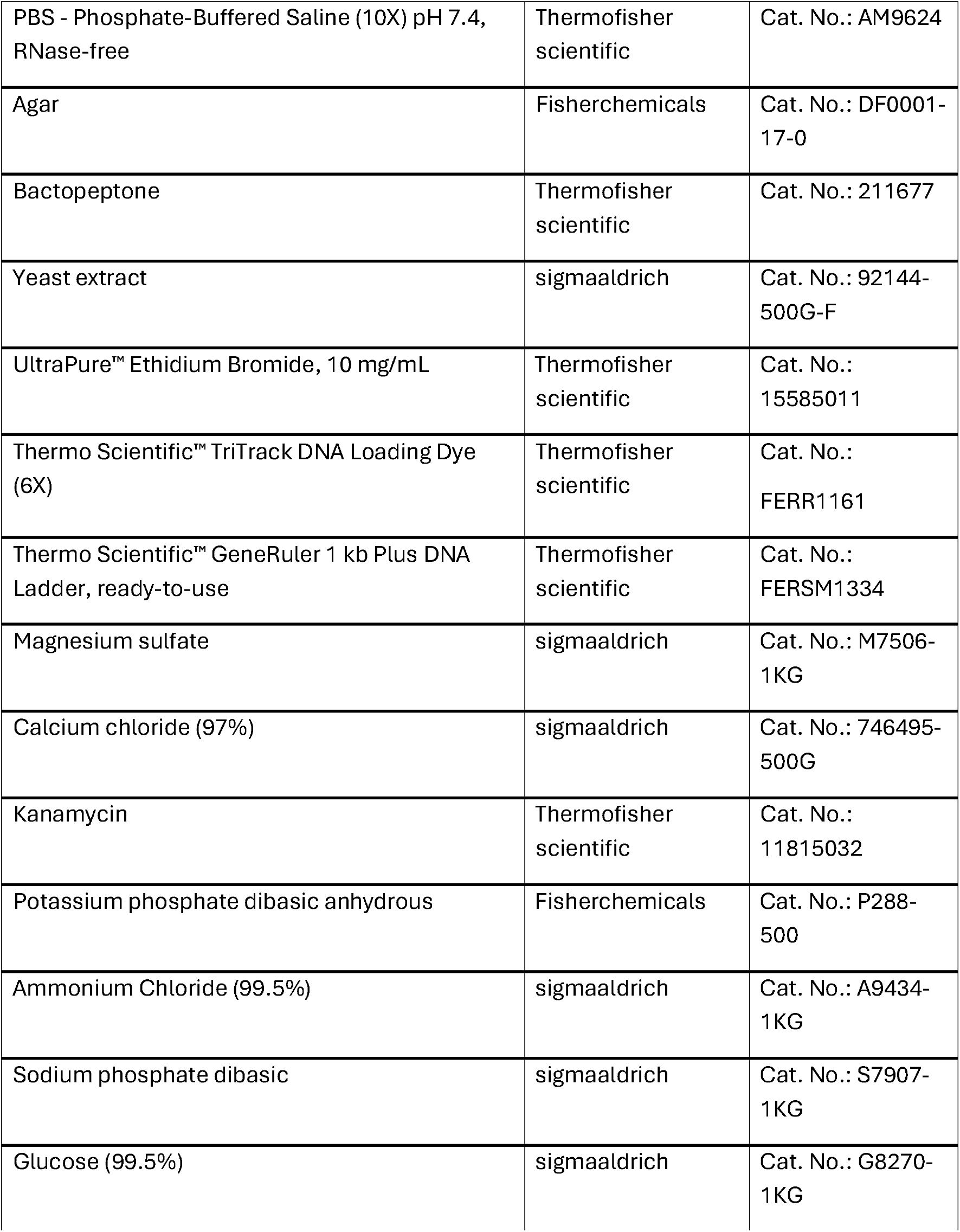
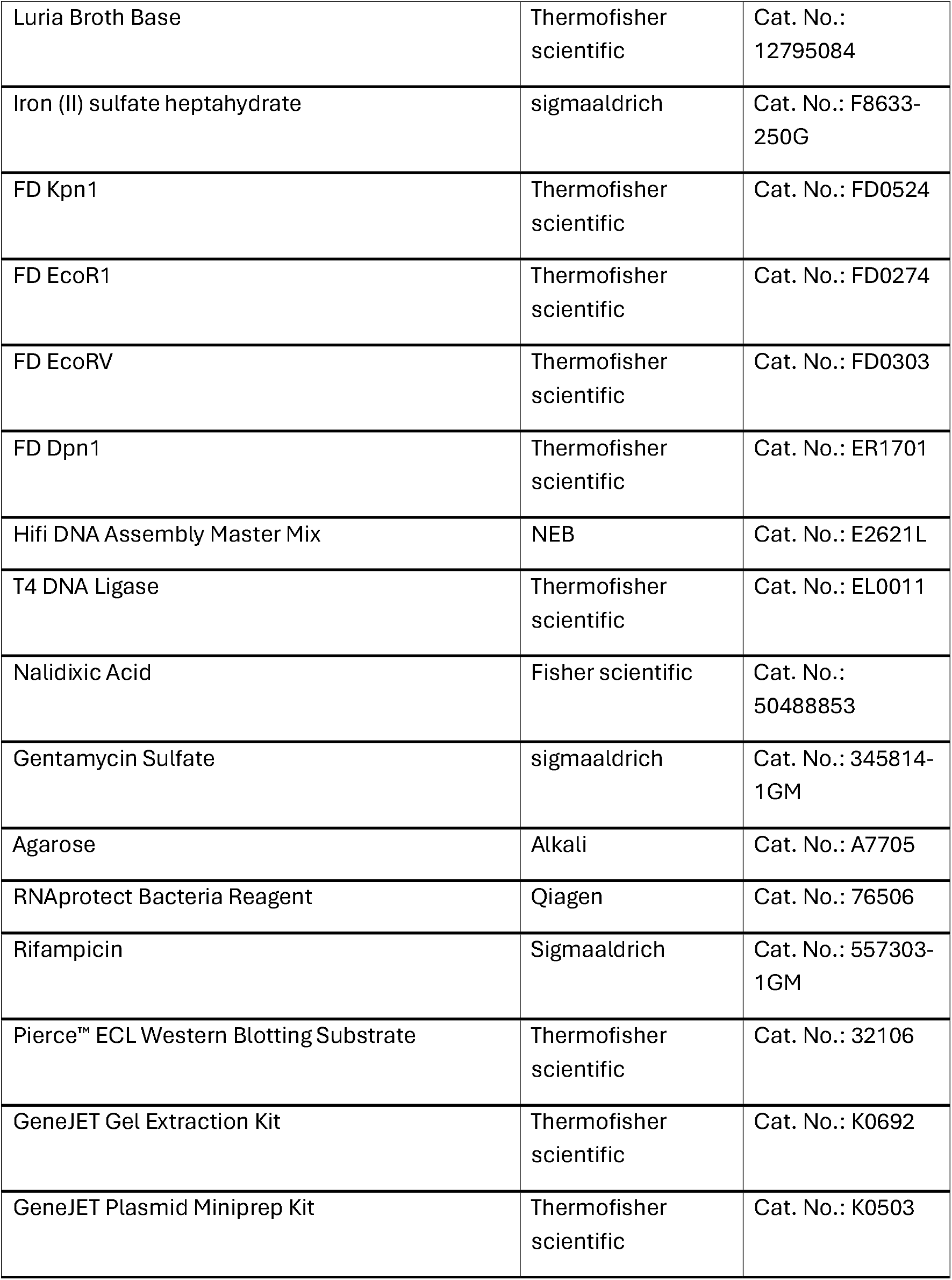
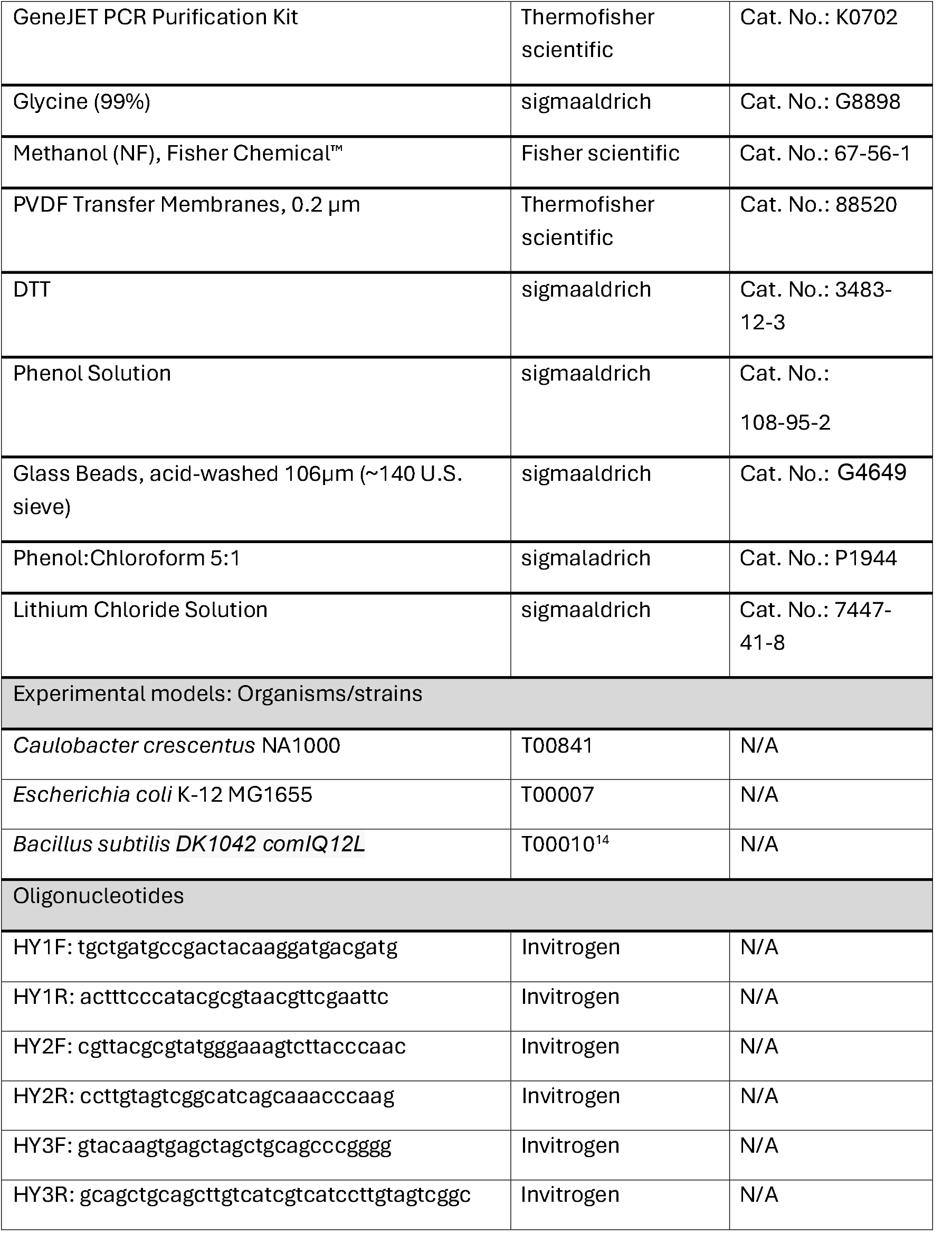
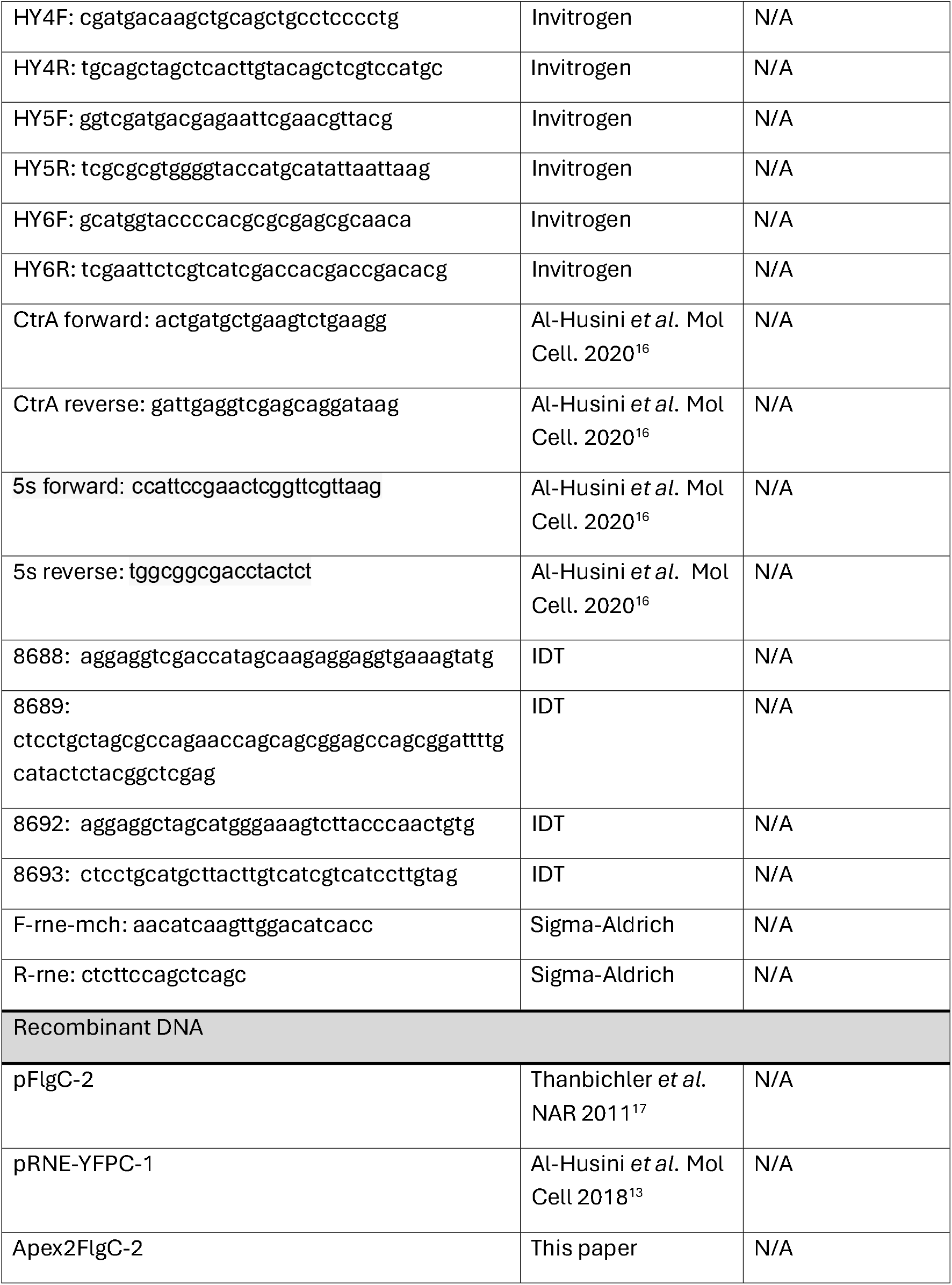
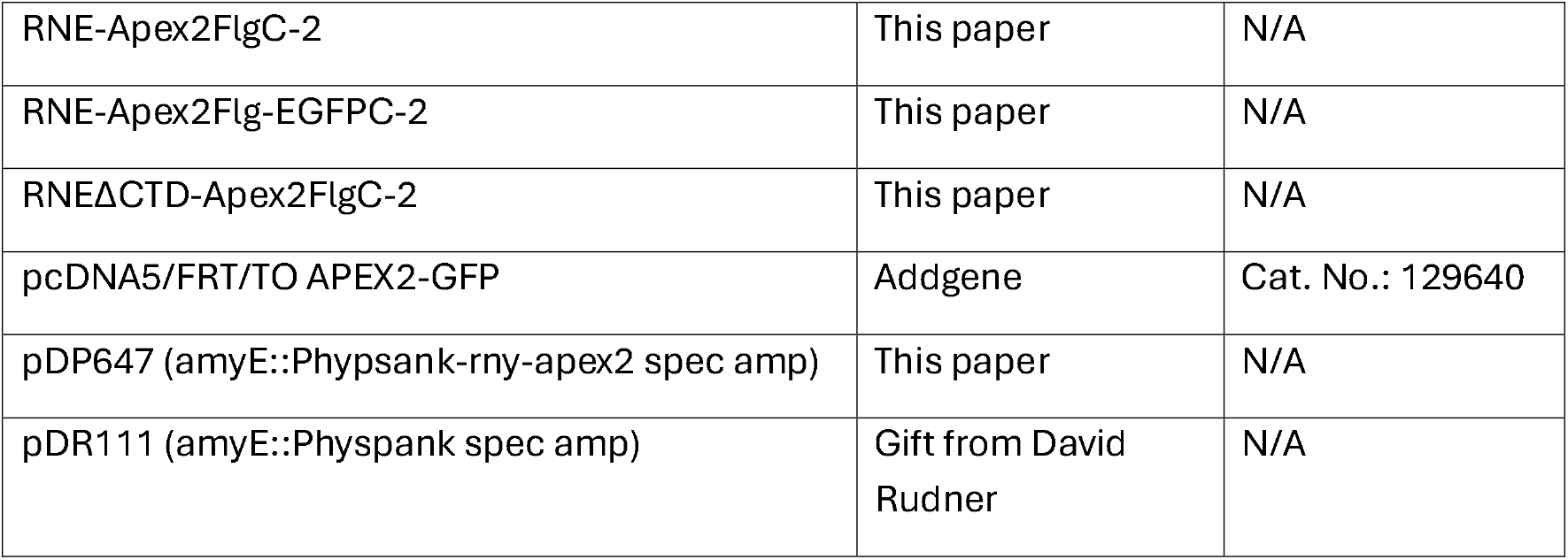
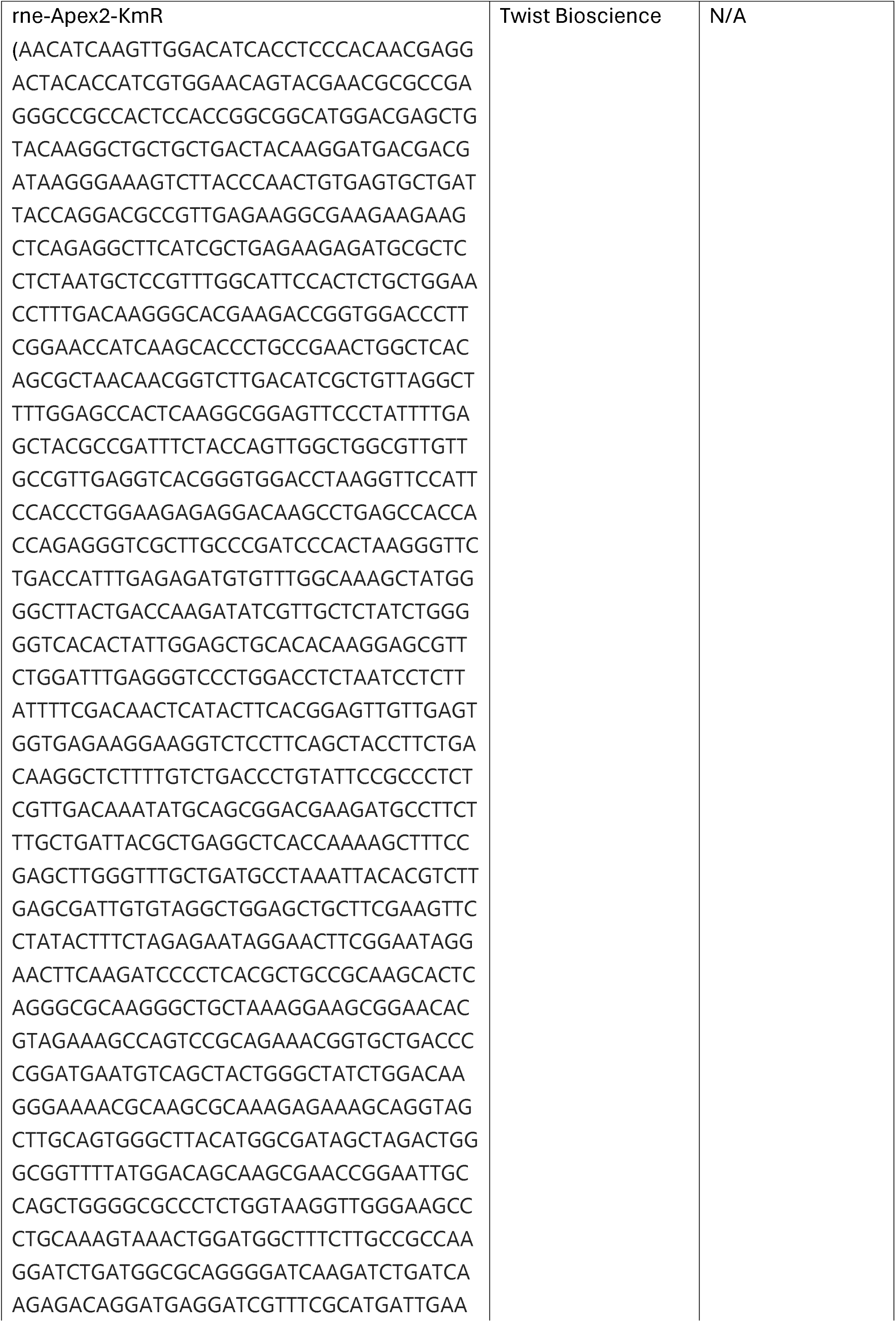

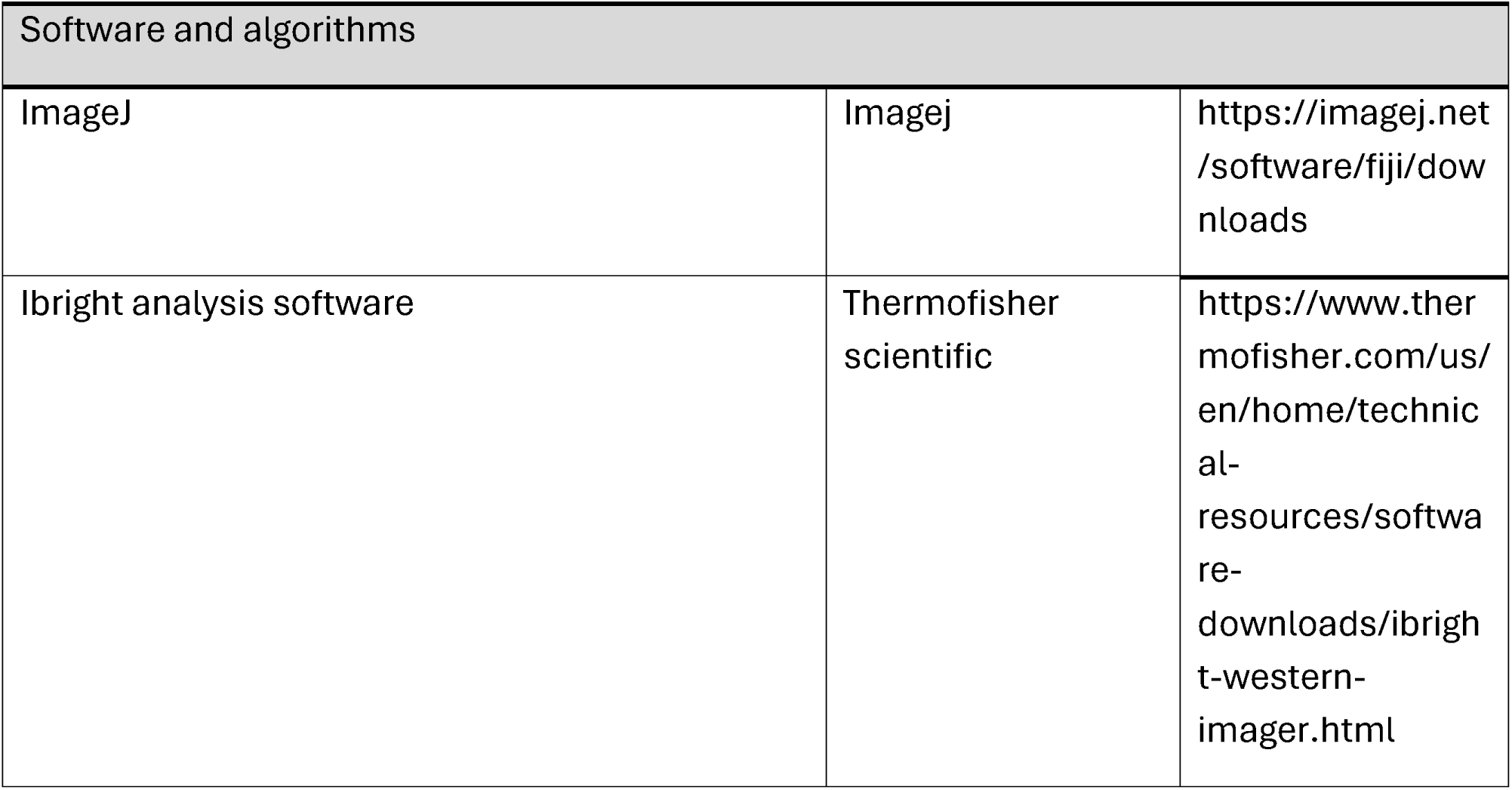

### *In vivo* RNA proximity labeling

#### APEX2, Alkyne-Phenol, and H_2_O_2_ are needed for RNA proximity labeling Gram negative bacteria

Gram negative bacteria were grown overnight in liquid media at their optimal growth temperature. Once the cultures reached an optical density (OD) of ∼0.3-0.6, 2.5 mM of alkyne-phenol was added and incubated for 30 minutes in a shaker incubator at 200 rpm. A 100 mM H_2_O_2_ solution (994 µL H_2_O + 6 µL H_2_O_2_, 50%) was then freshly prepared. 1 mM of H_2_O_2_ was then added for 45 seconds as the cultures were still shaking. The labeling experiment was carried out again with specific omissions as controls: no alkyne-phenol added, no H_2_O_2_ added, no alkyne-phenol and H_2_O_2_ added, different concentrations of alkyne-phenol added (0.5 mM and 1 mM), and different H_2_O_2_ exposures (15 seconds, 30 seconds, and 60 seconds). 4 mL of cell culture were then quickly transferred into two 2 mL Eppendorf tubes and centrifuged at 14,000 rpm for 10-15 seconds. The supernatant was quickly discarded, and the cell pellets were resuspended in 1 mL of 65°C pre-warmed TRIzol. Because proximity labeling requires the folded APEX2 protein, its denaturation in TRIzol can be used to quench the proximity labeling reactions. RNA extraction followed by ethanol precipitation was then carried out. 55 µL of a 10 mM Tris-HCl pH 7.0 (no EDTA) solution was added to dissolve the air-dried RNA pellets. The samples were vortexed briefly to aid resuspension. 5 µL were taken from the samples and placed in new tubes for Agilent Tapestation analyses and cell lysate RNA sequencing.

#### Gram positive bacteria

Gram positive bacteria were grown overnight in liquid media at their optimal growth temperature. Once the cultures reached an optical density (OD) of ∼0.3-0.6, the cultures were re-inoculated into fresh media at an OD of ∼0.05 supplemented with 1 mM isopropyl β-D-thiogalactopyranoside (IPTG) to induce the expression of RNY-APEX2. Once the cultures reached an OD of ∼0.4-0.5, 2.5 mM of alkyne-phenol was added and incubated for 30 minutes in a shaker incubator at 200 rpm. A 100 mM H_2_O_2_ solution was then freshly prepared.1 mM of H_2_O_2_ was then added for 45 seconds as the cultures were still shaking. An equal amount of cold methanol (pre-stored at -80 _°_C) was then added to the culture solution. The mixture was then quickly vortexed for 5 seconds and centrifuged at 14,000 rpm for 10-15 seconds. The cell pellets were resuspended in LETS buffer (1M Tris-HCl, 1M LiCl, 0.5M EDTA, and 10% SDS; for a 4 mL of gram positive cultures, the cell pellets were dissolved in 550 uL of LETS buffer). Both the LETS buffer and bead:phenol solutions were pre-warmed at 75 _°_C before being used. The solution was then transferred to a tube containing glass beads + phenol (for a 4 mL of gram positive culture, 320 uL of glass beads + 380 uL of phenol were used). The mixture was vortexed for 3 minutes. Chloroform was then added to the mixture and the solution was vortexed for 30 seconds (for a 4 mL of gram positive culture, 400 uL of chloroform was added to the cells + beads solution). The samples were then centrifuged at 3200 x g for 10 minutes. The top aqueous layer was then transferred to a fresh tube containing a phenol:chloroform solution (for a 4 mL of gram positive culture, 550 uL of phenol:chloroform was used). The phenol:chloroform solution was pre-warmed at 75 _°_C before being used. The mixture was then vortexed for 3 minutes, and the sample was centrifuged at 3200 x g for 10 minutes. The top aqueous layer was then transferred to a fresh tube and RNA precipitation was carried out. 55 µL of a 10 mM Tris-HCl pH 7.0 (no EDTA) solution was added to dissolve the air-dried RNA pellets. The samples were vortexed briefly to aid resuspension.

#### Copper-catalyzed click chemistry reactions

Copper-catalyzed click chemistry reactions were then conducted on the labeled RNA. The reactions have a 300 µL total volume and contained: 50 µL of 100 ng/uL RNA (5 µg) + 175 µL H_2_O + 12 µL of 125 mM sodium ascorbate + 3 µL of 10 mM Cy5-Azide + 60 µL of a mix of 2.5 mM Cu(II)SO_4_/12.5 mM THPTA (tris-hydroxypropyltriazolylmethylamine). The reagents were added in the order they are listed. The click reaction was vortexed for 5 seconds and incubated for 10 minutes at room temperature away from light exposure. Afterward, RNA precipitation was conducted and the RNA pellets were resuspended in 50 µL of 10 mM Tris-HCl pH 7.0 and 0.1 mM EDTA. The samples were then subjected to dot blot filtration on a nylon membrane following the manual’s instructions of BIO RAD’s Bio-Dot® and Bio-Dot SF Microfiltration Apparatus. A nylon membrane was pre-wetted in a 6X SSC solution for at least 10 minutes. The pre-wetted nylon membrane was then assembled into the dot blot apparatus. Vacuuming was applied to the apparatus and tighter sealing was implemented. The wells were then washed 3 times with 1 mL of an ice-cold mixture of 10 mM NaOH/1 mM EDTA. Afterward, the 5 µg RNA samples were directly applied to the wells (no RNA dissolution was performed). The wells were washed 3 times with 1 mL of an ice-cold mixture of 10 mM NaOH/1 mM EDTA. The blotted membrane was washed 3 times for 5 minutes with a 30 mL solution containing 2X SSC and 0.1% SDS. Fluorescent blot imaging was then visualized using an ibright imaging system at a fluorescent light exposure of 50 ms under the Cy5 channel.

#### RNase A & DNase I treatments

5 µg RNA (50 µL of 100 ng/µL of RNA) collected from a proximity labeling reaction (4 mL of Js767 cells grown in M2G media with 2.5 mM alkyne phenol, 45 seconds H_2_O_2_) were subjected to either an RNase A (5 µL of 10 mg/mL RNase A) or DNase I (5 µL of 1 U/µL DNase I) treatment. The treated and mock samples were incubated for 2 hours at 37°C. Following the enzyme or mock treatments, the RNA was precipitated and resuspended in 50 µL of 10 mM Tris-HCl pH 7.0 and then subjected to the Copper catalyzed click-chemistry reaction with Cy5-azide. Following the 10-minute click reaction incubation, the RNA was then precipitated and subjected to dot blot filtration as previously described.

#### Streptavidin capture of biotin labeled RNA

To enrich the transcriptome of BR-bodies, RNA labeled with alkyne-phenol was subjected to the following click chemistry reaction: 50 µL of RNA (total RNA volume) + 175 µL H2O + 12 µL of 125 mM sodium ascorbate + 3 µL of 10 mM Biotin-peg3-azide + 60 µL of a mix of 2.5 mM Cu(II)SO_4_/12.5 mM THPTA. The reagents were added in order as they are listed. The click reaction was vortexed briefly and incubated for 10 minutes at room temperature away from light exposure. RNA precipitation was then conducted and the air-dried RNA pellets were resuspended in 100 µL B&W buffer (5 mM Tris-HCl, pH 7.5, 1 M NaCl, 0.5 mM EDTA, 0.1% v/v tween-20). The RNA samples were then immediately used for biotinylated RNA enrichment.

High-capacity magnetic streptavidin beads (Vectorslab) were washed three times with 1 mL B&W buffer, twice with 1 mL of a mix of 0.1 M NaOH/0.05 M NaCl, and once with 1 mL 0.1 M NaCl at room temperature. The beads were then blocked using a blocking buffer (1 mg/mL BSA + 1 mg/ml heparin salt, dissolved in B&W buffer) at room temperature for 2 hours. The beads were then washed three times with 1 mL of B&W buffer under gently vortexing. Afterward, the 100 µL RNA solution was added to the beads along with an extra 100 µL of B&W buffer. The RNA and beads were allowed to mix at room temperature for one hour. The RNA-loaded beads were washed three times with 1 mL of B&W buffer, twice with 1 mL of a PBS solution containing 4 M urea + 0.1% SDS, and twice with 1 mL of just PBS at room temperature. To elute the enriched RNA, 900 uL of 65°C pre-warmed TRIzol was added to the beads and the samples were incubated at 65°C for 10 minutes. Afterward, 200 µL of chloroform was added and the samples were inverted 6-7 times and then were incubated for 5 minutes at room temperature. The samples were centrifuged at 14,000 rpm for 10 minutes at 4°C. The upper aqueous layer was transferred to a new Eppendorf tube and the enriched RNA was precipitated as previously described. The air-dried RNA pellets were resuspended in 10 µL of 10 mM Tris-HCl pH 7.0 and 0.1 mM EDTA. 4 µL were then used for Agilent Tapestation analysis. The remaining volume could be utilized for the follow-up RNA-seq experiments.

#### RNA quality assessment

Js767 (RNE-Apex2FlgC-2) was grown in PYE at 28°C overnight and re-inoculated into 5 mL M2G the next day. Once the cultures reached an OD of 1.2, 2.5 mM of alkyne-phenol was added and incubated for 30 minutes in a shaker incubator at 28°C at 200 rpm. 1 mM of H_2_O_2_ was added to the culture for 45 seconds. Total RNA was then collected and precipitated from 4 mL of culture as previously described using TRIzol. Afterward, 50 µL of 100 ng/µL of alkylated RNA were subjected to the following copper catalyzed click-chemistry reaction: 50 µL of 100 ng/µL RNA (5 µg) + 175 µL H_2_O + 12 µL of 125 mM sodium ascorbate + 3 µL of 10 mM Cy5-Azide + 15 µL of 100 mM aminoguanidine hydrochloric acid + 60 µL of a mix of 2.5 mM Cu(II)SO_4_/12.5 mM THPTA. The click reaction was incubated for 10 minutes at room temperature away from light exposure. The RNA from the click reaction was then precipitated and the air-dried RNA pellet was resuspended in 50 µL of 10 mM Tris-HCl pH 7.0 and 0.1 mM EDTA. 250 ng of RNA from the Js767 RNA lysate and the click reaction RNA were sent for Agilent Tapestation analysis.

### RNA precipitation

RNA precipitation was carried out by adding, in order, 2 µL of 1 mg/mL glycogen, 1/10 sample volume of 3 M sodium acetate pH 5.5, and 1 sample volume of isopropanol. The sample mixtures were vigorously vortexed and stored at -80°C overnight. The samples were spun for 1 hour at 14,000 rpm. The supernatant was then decanted and 1 mL of ice-cold 80% ethanol was added. The samples were spun for 15 minutes at 14,000 rpm. The ethanol was pipetted out and the samples were then spun for 1 minute at 14,000 rpm. Any remaining ethanol was pipetted out and the pellets were allowed to air-dry until they became translucent. The RNA was then resuspended in the buffer of choice.

### Western Blot

The following buffers were initially prepared:

1. 10X transfering buffer: 144 grams of glycine and 30.2 grams of Tris-Base were dissolved in 900 mL of ddH2O. The buffer volume was adjusted to 1 L of ddH2O and then filtered utilizing a Nalgene Rapid-flow sterile disposable filter unit.
2. 10X TBS: 24 grams of Tris-Base and 88 grams of NaCl were dissolved in 900 mL of ddH2O. The pH of the buffer was adjusted to 7.6 utilizing hydrochloric acid. The buffer volume was adjusted to 1 L of ddH2O and then sterilized utilizing a Nalgene Rapid-flow sterile disposable filter unit.
3. 1X TBST: 10X TBS was diluted 10-fold with ddH2O to make 1X TBS. 0.1% Tween-20 was then added to the 1X TBS solution to make 1X TBST.

NA1000, LS4379 (hfq-m2), and Js767(RNE-Apex2FlgC-2) were inoculated in PYE and incubated overnight at 28°C until they reached log phase (OD ∼0.3-0.6). 1 mL was taken from each culture, placed in 1.5 mL Eppendorf tubes, and centrifuged for 2 minutes at 6,000 rpm. The liquid cultures were discarded, and the cells were resuspended in 4X SDS loading dye. 125 µL of 4X SDS loading dye was added to cultures with an OD∼0.5. The samples were boiled at 95°C for 5 minutes then vortexed vigorously. The vortexed samples were quickly spun down and placed on ice. 5 µL of a pre-stained PageRuler marker and 20 µL of lysate from each sample were loaded onto an SDS-PAGE gel. 1X transferring buffer was freshly prepared (20 mL of 10X transfer buffer + 20 mL of methanol in 160 mL ddH2O) while the samples were running on the gel. A PVDF transferring membrane was wetted in methanol and 6 blotting papers were wetted using 1X transferring buffer for at least 10 minutes. Once the lysate samples on the SDS-PAGE were properly resolved, the gel was placed on top of 3 pre-wetted blotting papers, and 2 mL of 1X transferring buffer was poured on the gel. The PVDF membrane was briefly plunged into the 1X transferring buffer and then placed on top of the gel. The remaining 3 blotting papers were placed on top of the PVDF membrane. The transferring of the lysates from the SDS-PAGE gel to the PVDF membrane occurred by utilizing the BIO-RAD Trans-Blot Turbo Transfer system (1 amps, 2.5 volts, 15 minutes). Once the transfer was completed, the PVDF membrane was placed in a new container containing 5% BSA (blocking solution). The container was nutated for 1 hour at room temperature. Afterward, the 5% BSA blocking solution was discarded and the PVDF membrane was washed 5 times with ddH2O for 5 minutes per wash. The primary antibody (DYKDDDDK anti-flag tag) was then diluted (1:10000) into a fresh 5% BSA solution. Once the last ddH2O wash was completed, the primary antibody solution was poured on the PVDF membrane, and the container was nutated at 4°C overnight. The following day, the primary antibody solution was discarded and the PVDF membrane was washed 5 times with TBST for 5 minutes per wash. The secondary antibody (goat anti-rabbit secondary antibody, HRP conjugated) was then diluted (1:10000) into a fresh 5% BSA solution. Once the last TBST wash was completed, the secondary antibody solution was poured on the PVDF membrane, and the container was nutated at room temperature for 1 hour avoiding light exposure. The secondary antibody solution was then discarded and the PVDF membrane was washed 5 times with TBST for 5 minutes per wash. Following the final washing procedure, a pierce ECL western blotting substrate solution was prepared (1 mL of reagent 1 + 1 mL of reagent 2 were mixed and vigorously vortexed). The PVDF membrane was placed on Seram wrap and the 2 mL substrate solution was poured on top of the membrane. The substrate was incubated on the PVDF membrane at room temperature for 5 minutes avoiding light exposure. The membrane was then placed into an iBright imaging system and the signal was detected under the Chemi-blot filter.

### Dilution Plates

NA1000 & Js767 (RNE-Apex2FlgC-2) were grown in PYE overnight at 28°C. The overnight cultures were diluted into fresh media and incubated at 28°C until the cells reached an OD of ∼0.3-0.6. Cells were then diluted to an OD=0.05 in PYE and 4 10-fold serial dilutions were made. 5 µL were spotted from each dilution onto PYE plates. The plates were incubated at 28°C for two days, and images were taken using an ibright imaging system.

### Fluorescent Cell Imaging

Js87 (RNE-msfGFP) & Js768 (RNE-Apex2FlgC-EGFPC-2) were grown in PYE + gent (0.5 µg/mL) and PYE + Kan, respectively, overnight at 28°C. Cells were then fixed on M2G + 1.5% agarose pads placed on microscope slides (3051, Thermofisher scientific). Imaging was performed on an epifluorescence microscope with a 100X objective. Specifically, Nikon elements software was used to control a Nikon Eclipse NI-E equipped with a CoolSNAP MYO-CCD camera and a 100x Oil CFI Plan Fluor (Nikon) objective to capture the images.

### mRNA half-lives measurement

NA1000 and Js767 (RNE-Apex2FlgC-2) were grown in liquid PYE at 28°C overnight. Cells were then serially re-inoculated into liquid M2G and incubated until overnight log phase (OD ∼0.3-0.6) was reached. Log phase cultures were then re-inoculated to an OD = 0.05 in 25 mL liquid M2G. Before adding rifampicin, at time point 0, 1 mL of cells were added to 2 mL of RNAprotect Bacterial reagent (Qiagen) and vortexed for 5 seconds. 200 µg/mL of Rifampicin was administered to the cultures and 1 mL of cells were extracted and added to 2 mL of RNAprotect Bacterial reagent at each of the following time points (followed by 5 seconds vortexing): 1, 2, 4, and 8 minutes. Cells were incubated at room temperature in the RNAprotect Bacterial reagent for 5 minutes before being spun at 5000 rpm for 5 minutes. The bacterial pellets were resuspended in 1 mL of 65°C pre-heated TRizol (Ambion) and incubated at 65°C for 10 minutes. 200 µL of chloroform was then added and the samples were incubated at room temperature for 5 minutes. Subsequently, the samples were centrifuged at 14,000 rpm for 10 minutes at 4°C. The aqueous layer was removed and placed in a new 1.5 mL Eppendorf tube. The RNA was then precipitated and the pellets were resuspended in 50 µL elution buffer (10 mM Tris-HCl, pH=7.0, 0.1 mM EDTA). PCR tubes were filled with a master mix that contained 0.4 µM of ctrA forward primer and 0.4 µM of ctrA reverse primer, 1X Luna Universal One-Step Reaction Mix, 1X Luna WarmStart RT Enzyme Mix, and water. 100 ng/µL of RNA template was additionally aliquoted into each of the PCR tubes. The samples were mixed well and quickly spun down. A QuantStudio Real-Time PCR apparatus was utilized to conduct and examine the qRT-PCR experiments. The same qRT-PCR experiment was done using instead the 5S forward and reverse primers. To determine the mRNA-decay rates, we fitted a linear curve to the ln (fraction RNA remaining) at each time point. Using a standard curve, the Ct was converted into the quantity of RNA. Each time point’s 5S rRNA amount was divided by the amount of 5S rRNA at time point 0. The number obtained at each time point was then used to normalize the amount of ctrA at each time point. The natural log of % RNA remaining found in each sample was divided by the natural log of RNA at time point 0. The slopes of the linear curve fit were then converted into mRNA half-life using the following equation: mRNA half-life=−ln(2)/slope.

### RNA-seq

Duplicate biological samples were prepared for RNA-seq of the Full-length RNase E-APEX2 fusion (JS767) and the RNase E-NTD-APEX2 (JS801). Lysates from each sample, as well as the eluted RNA from affinity purification were prepared for RNA-seq. RNA-seq was performed by IU-Bloomington CGB genomics service facility using the Illumina Tru-seq stranded HT kit. Raw sequencing reads were deposited to the NCBI GEO database with accession number GSE297938. Data analysis was performed similar to Al-Husini et al. Mol Cell 2020^16^. RNA reads were stripped of adapters, and tRNA and rRNA reads were aligned to unique file with duplicate copies deleted using bowtie^18^, and reads not aligning to tRNA and rRNA were aligned to the C. crescentus transcriptional units^19^ using bowtie. Genes with fewer than 50 reads mapping to them were omitted from analysis, and the RPKM values were calculated for each sample. The log2 ratio between the RPKM values in the elution and lysate samples were then calculated on all genes with >=50 reads in both elution and lysate samples. Samples with log2(elution/lysate) measurements in all four biological samples were used for comparison for BR-body proximity labeling enrichment. To calculate which RNA types were enriched in BR-bodies, the distribution of enrichment measurements log2(elution RPKM/lysate RPKM) were subjected to a T-test with unequal variance between the BR-body + (JS767) and BR-body – (JS801) strains and the resulting p-values were reported. Raw read count and processed proximity labeling RNA-seq data can be found in Table S3.

### Plasmid Construction

#### pAPEX2-FlgC-2 *Kan*^*R*^

The pFlgC-2 vector^17^ was PCR amplified using primers HY1F & HY1R. APEX2 was PCR amplified using primers HY2F & HY2R from the addgene template #129640. The amplicons were run on a 1% agarose gel and gel extracted using a GeneJET Gel Extraction Kit. The purified vector was Dpn1 treated then column purified using the GeneJET PCR Purification Kit. The Apex2FlgC-2 *Kan*^*R*^ plasmid was assembled via Gibson assembly (NEB) and transformed into chemically competent DHbeta10 E. coli and plated on LB + Kan (50 µg/mL) plates. The resulting KanR colonies were minipreped using the GeneJET Plasmid Miniprep Kit and screened via restriction digestion (EcoR1) and the insert sequence was verified by Sanger sequencing (Genewiz).

#### pRNE-Apex2-FlgC-2 *Kan*^*R*^

The last 534 RNase E (RNE) base pairs were obtained by digesting pRNE-YFPC-1^17^ with Nde1 and Kpn1. APEX2FlgC-2 was digested with Nde1 and Kpn1. The digestion reactions were run on a 1% agarose gel and the RNE fragment and Apex2-FlgC-2 vector were gel extracted using a GeneJET Gel Extraction Kit. The RNE fragment and Apex2FlgC-2 vector were ligated using T4 ligase, transformed into chemically competent DHbeta10 E. coli, and selected on LB + Kan plates. The resulting KanR colonies were minipreped using the GeneJET Plasmid Miniprep Kit and screened via restriction digestion (EcoRV) and the insert sequence was verified by Sanger sequencing (Genewiz).

#### pRNE-Apex2FlgC-EGFPC-2 *Kan*^*R*^

The RNE-Apex2FlgC-2 vector was PCR amplified using primers HY3F & HY3R. The EGFP insert was PCR amplified using primers HY4F & HY4R from the addgene template #129640. The amplicons were run on a 1% agarose gel and gel extracted using a GeneJET Gel Extraction Kit. The purified vector was Dpn1 treated then column purified using the GeneJET PCR Purification Kit. The RNE-Apex2FlgC-EGFPC-2 plasmid was assembled via Gibson assembly (NEB) and transformed into chemically competent DHbeta10 E. coli and selected on LB + Kan plates. The resulting KanR colonies were minipreped using the GeneJET Plasmid Miniprep Kit and screened via restriction digestion (EcoRV) and the insert sequence was verified by Sanger sequencing (Genewiz).

#### pRNEΔCTD-Apex2FlgC-2 Kan^R^

The pApex2-FlgC-2 vector^17^ was PCR amplified using primers HY5F & HY5R. RNE-NTD was PCR amplified using primers HY6F & HY6R from NA1000 cells. The amplicons were run on a 1% agarose gel and gel extracted using a GeneJET Gel Extraction Kit. The purified vector was Dpn1 treated then column purified using the GeneJET PCR Purification Kit. The pRNE(NTD)-Apex2FlgC-2 *Kan*^*R*^ plasmid was assembled via Gibson assembly (NEB) and transformed into chemically competent DHbeta10 E. coli and plated on LB + Kan (50 µg/mL) plates. The resulting KanR colonies were minipreped using the GeneJET Plasmid Miniprep Kit and screened via restriction digestion (Kpn1) and the insert sequence was verified by Sanger sequencing (Genewiz).

#### pDP647 amyE::Phypsank-rny-apex2 spec^R^ amp^R^

To generate the B. subtilis inducible Rny-APEX2 construct pDP647, the rny gene was amplified from wild type DK1042 DNA with primers 8688/8689 and apex2 was amplified from plasmid pAPEXC-2 with primers 8692/8693. Next, the rny amplicon was digested with SalI and NheI, the apex2 amplicon was digested with NheI and SphI and the two fragments were simultaneously ligated into the SalI and SphI sites of pDR111 that carries a polylinker downstream of the IPTG-inducible P_hyspank_ promoter, the gene encoding the LacI repressor, and a spectinomycin resistance cassette between the arms of the amyE gene (generous gift of David Rudner, Harvard Medical School).

### Strain construction

#### Js767: NA1000 rne::rne-apex2-flg Kan^R^

The RNE-Apex2FlgC2 plasmid was recombined into the rne locus in NA1000 via mating and the selection was carried out on PYE + Nal (20 µg/mL) + Kan (25 µg/ mL) plates. The resulting KanR colonies were first grown in PYE + Kan (5 µg/mL) cultures and then screened by PCR.

#### Js801: NA1000 rne::rneΔCTD-apex2-flg Kan^R^

The RNEΔCTD-Apex2FlgC2 plasmid was recombined into the rne locus in NA1000 via mating and the selection was carried out on PYE + Nal (20 µg/mL) + Kan (25 µg/ mL) plates. The resulting KanR colonies were first grown in PYE + Kan (5 µg/mL) cultures and then screened by PCR.

#### Js768: NA1000 rne::rne-apex2-flg-egfp Kan^R^

The RNE-Apex2FlgC-EGFPC-2 plasmid was recombined into the rne locus in NA1000 via mating and the selection was carried out on PYE + Kan plates. The resulting KanR colonies were first grown in PYE + Kan cultures and then screened by PCR.

#### DB2579 amyE::rny-APEX2 spec^R^ amp^R^

The pDP647 plasmid was transformed into DK1042^14^ cells, and selected on plates containing LB and 100 µg/mL Spectinomycin. The resulting colonies were then screened for the integration by PCR.

#### ES413: rne::rne-mcherry-flg-Apex2 kan^R^

The in vitro synthesized sequence rne-Apex2-KmR was PCR amplified using F-rne-mch and R-rne primers and recombined into the rne locus in Kti162^15^ by lambda red system^20^, the selection was carried out on LB + Kan (30 µg/ mL) plates. The rne-mcherry-flag-Apex2 kan^R^ was then moved to MG1655 by P1 transduction^21^. The resulting Kan^R^ colonies were grown in LB + Kan, screened by PCR and verified by Sanger sequencing (The Center for Genomic Technologies – Huji).

## Results

### RNaseE-APEX2 proximity labeling of RNA requires Alkyne-Phenol, H_2_0_2_, and APEX2

APEX2 is an engineered ascorbate peroxidase which can catalyze the creation of a free radical on the oxygen of alkyne-phenol, and this highly reactive species can react with molecules within a 10-20 nm radius^22,23^ (Fig 1A). It was shown that APEX2 can label RNA in eukaryotic cells, making this a useful experimental system for identifying localized RNAs^10,12,24,25^. In order to perform RNA proximity labeling in bacteria, we first generated a gene fusion between RNase E and APEX2. RNase E is known to phase-separate into bacterial ribonucleoprotein bodies (BR-bodies), biomolecular condensates containing mRNA and promoting the mRNA decay process^13,16^. To determine whether APEX2 fusions are tolerated in bacteria, we examined the subcellular localization, cell fitness, and mRNA decay activity of RNase E-APEX2 fusions in the bacterium Caulobacter crescentus (Fig 1B). We observed that fusing APEX2 to RNase E led to proper expression of RNase E-APEX2 (Fig S1) and did not alter its ability to phase separate into BR-bodies (Fig 1B). In addition, since RNase E’s ability to degrade mRNAs is essential for cell growth^16,26^, we also examined the cellular fitness of the RNase E-APEX2 fusion and found that both CFUs and colony size were indistinguishable from wild-type (Fig 1B). Finally, we compared the mRNA decay activity of the RNase E-APEX2 fusion and found that it has similar rates of mRNA decay as compared to wild-type (Fig 1B), while 5S rRNA remained stable. Altogether, we could not detect any measurable differences in RNase E function when fused to APEX2. To examine whether the APEX2 fusion can be used to label RNA we added all combinations of proximity labeling reactants (H_2_O_2_ and Alkyne-Phenol) in combination with or without the RNase E-APEX2 fusion (Fig 1C). Each of the proximity labeling conditions were performed, and then we extracted the RNA and used copper click chemistry to conjugate a Cy5-azide to the RNA. The RNA was then deposited on a positively charged nylon membrane using a dot blot apparatus, and the Cy5 signal was measured in a fluorescent gel imager. We find that only in the presence of APEX2, H_2_O_2_, and alkyne-phenol do we observe RNA labeling (Fig 1C), suggesting that APEX2 proximity labeling reactions occurred. To determine whether the Cy5 signal was a result of RNA, or contaminating DNA we performed RNase A or DNase I digestions on our samples before spotting them on the dot blot (Fig 1C). Here we see that RNase A treatment leads to a complete loss of Cy5-fluorescence, while DNase I treatment led to no difference in fluorescence signal, suggesting that our Cy5 fluorescence is from RNA.

**Figure 1.**
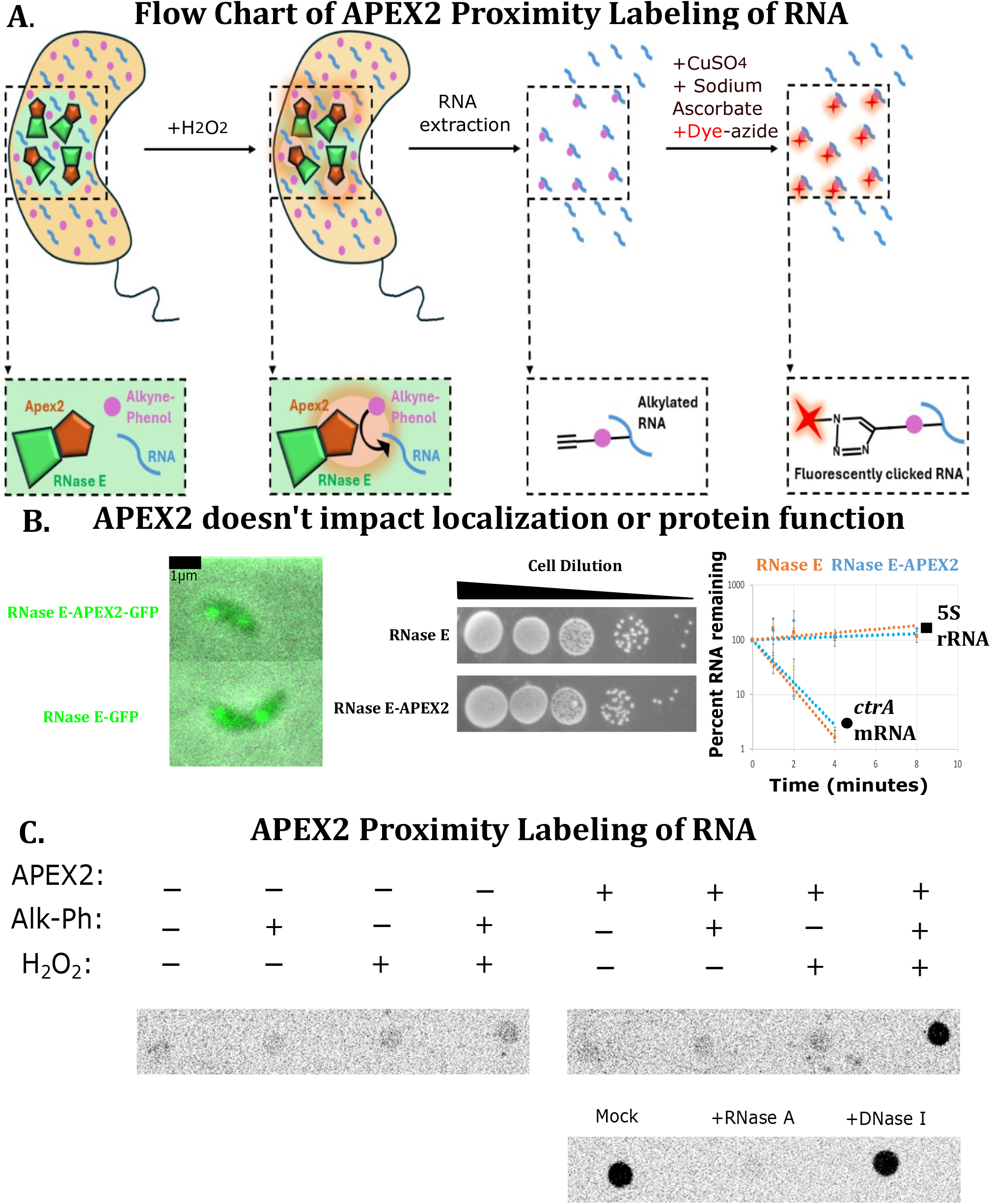
APEX2 proximity labeling of RNA in bacterial cells with minimal perturbation. A) Schematic of APEX2 labeling of RNA protocol. The APEX2 protein was fused to RNase E, the major protein that scaffolds BR-bodies. Cells are incubated in media containing alkyne-phenol, and labeled was initiated with H_2_O_2_. After a brief labeling reaction, RNA was extracted from the cells, and Cy5-azide was conjugated to the RNA by copper catalyzed click chemistry. B) APEX2 fusion does not dramatically impact localization or function (growth rate). Left: In vivo localization of RNase E-msfGFP vs RNase E APEX2-msfGFP shows that fusion does not impact the formation of BR-bodies. Scale bar is 1µm. Middle: Growth of RNase E-APEX2 fusion is similar to wild type Caulobacter cells. Right: mRNA half-life measurements by qRT-PCR show that RNase E-APEX2 degrades mRNAs with a similar half-life to wild-type. Data are from three biological and technical replicates. C) APEX2 labeling requires H_2_O_2_ and alkyne-phenol to label RNAs. RNA labeling reactions were placed on a nylon membrane to bind to the RNA in a dot blot apparatus and scanned for Cy5 fluorescence in a gel imager. As a control, the RNA samples were incubated for 2 hours with DNase I and RNase A. The RNA was then precipitated before being subjected to the azide-Cy5 click chemistry reaction and was re-precipitated before being filtered on the nylon membrane in the dot blot apparatus.

### Rapid labeling of cellular RNA with Alkyne-Phenol

One of the major challenges of studying bacterial mRNA decay is the very short lifetimes of cellular mRNAs. Most bacterial mRNAs in rapidly growing species have half-lives between 1-4 minutes^27,28^, making it hard to harvest the RNA before they are degraded. To address this technical limitation, we first optimized the Alkyne-phenol concentration used for labeling cells and found that 2.5 mM Alkyne-Phenol robustly labels cellular RNA (Fig 2). Next, we performed a time-course of H_2_O_2_ exposure to identify the timescale of RNA labeling, ranging from 15 seconds reaction times to 1 minute reaction times. We found that robust labeling could be achieved in as little as 15 seconds, while labeling increased upon longer proximity labeling reaction times (Fig 2). Importantly, the labeling reaction occurs on the sub-minutes timescale, making this method well-suited to the use in exponentially growing bacterial cells.

**Figure 2.**
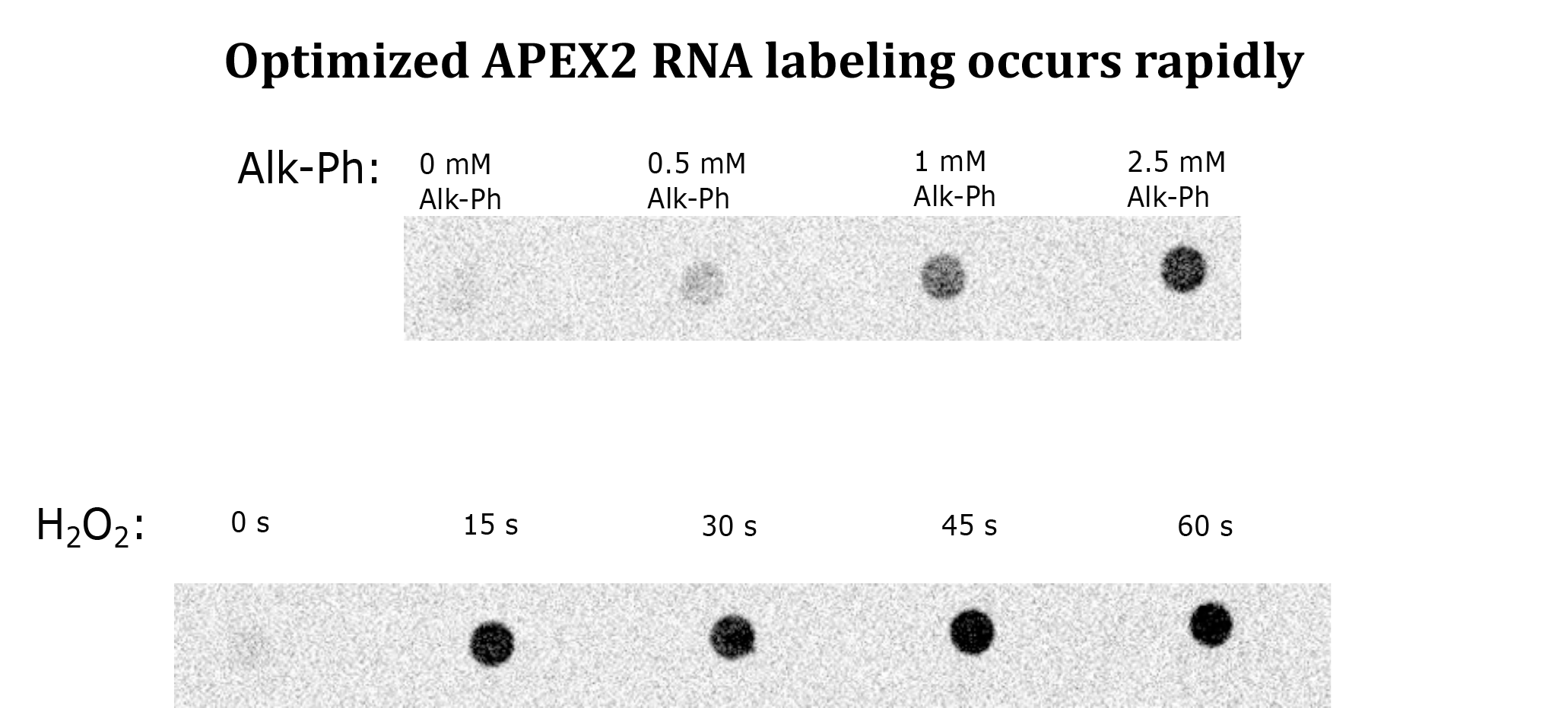
APEX2 proximity labeling of RNA works rapidly. Optimization of APEX2 labeling of RNA with Alkyne-Phenol. Top: Alkyne-phenol titration reveals that peak labeling occurs with 2.5 mM Alkyne-Phenol. RNA was labeled in the scheme shown in Fig 1A, and the Cy5 intensity was measured in a gel imager. Bottom: Time course of APEX2 labeling. RNA was labeled in the scheme shown in Fig 1A, and RNA labeling is apparent in as short as 15 seconds of H_2_O_2_ incubation, while peak labeling efficiency is observed at 45 seconds of H_2_O_2_ incubation.

### RNA proximity labeling works across bacteria

To increase the applicability of this method, we sought to determine whether APEX2 can label RNA in other species of bacteria. As a key requirement of APEX2 activity is heme, we chose to fuse APEX2 to RNA degradosome scaffold proteins in E. coli (gram -) and B. subtilis (gram +) as both organisms have heme biosynthesis pathways encoded in their genomes. In E. coli, we fused APEX2 to RNase E, which scaffolds BR-bodies in this species^15,29^, localizing into membrane anchored RNP foci^15^. In B. subtilis, we fused APEX2 to RNase Y, which scaffolds BR-bodies in this species and also localizes into RNP foci on the inner membrane of the cell^29,30^. In both of these species, we observed RNA labeling after 45 seconds of H_2_O_2_ incubation that required the APEX2 fusion (Fig 3), suggesting that APEX2 proximity labeling of RNA can be used in diverse species of bacteria which encode heme biosynthesis pathways.

**Figure 3.**
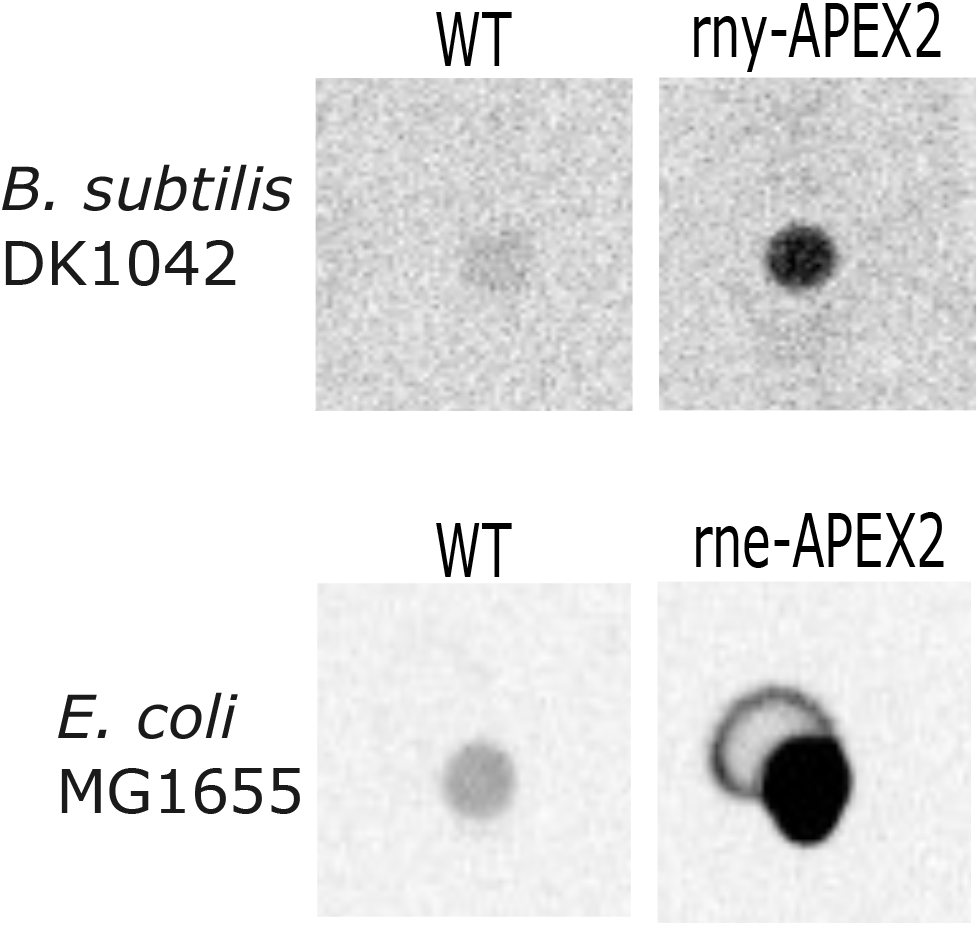
APEX2 proximity labeling of RNA works across species. APEX2 requires heme for activity, so APEX2 fusions were generated in two additional species with heme biosynthesis pathways, B. subtilis (gram +) and E. coli (gram –). Both species were and subjected to RNA proximity labeling in mid-exponential phase of growth in LB media at 37°C, pre-incubated with alkyne-phenol for 30 minutes, and proximity labeling was triggered by a 45 second incubation with H_2_O_2_. RNA labeling reactions were subjected to azide-Cy5 click chemistry reactions, and 5 μg RNA placed on a nylon membrane to bind to the RNA in a dot blot apparatus and scanned for Cy5 fluorescence in a gel imager.

### Copper click-chemistry of Azide-Biotin and streptavidin purification of labeled RNA

APEX2 proximity labeling of RNA is rather flexible due to the diversity of azides commercially available for copper catalyzed click chemistry. To isolate cellular RNA for RNA-sequencing, we altered the azide from azide-Cy5 dye to azide-biotin to allow for streptavidin-mediated RNA purification from the cell (Fig 4A). In this approach, labeled RNA is extracted from cells and conjugated to azide-biotin using copper catalyzed click chemistry, which allows for the subsequent purification of the labeled RNA using streptavidin-resin. While copper can cleave RNA, using a short incubation time for the click reaction minimizes RNA cleavage by copper (Fig S2). After click chemistry, the RNA can be purified under stringent conditions and eluted from the resin under denaturing conditions. When applied to the RNase E-APEX2 fusion presented earlier, we found that the eluted RNA profile indeed matched that of BR-body RNA isolated via density centrifugation (Fig 4B)^16,31^. To compare the specificity of APEX RNA proximity labeling, we prepared eluted RNAs from BR-body + cells (JS767) and BR-Body – cells (RNaseEΔCTD-APEX2, JS801) and prepared them for RNA-seq. As a control for the initial amount of each RNA in the cell, we also performed RNA-seq on the total RNA lysates of each strain. The overall level of proximity labeling was calculated as the log2 enrichment (eluted/lysate) of each RNA measured at >50 reads/sample. Overall, we found that BR-body + samples (JS767) contained higher overall enrichment levels than BR-body – (JS801) (Fig S3A) whose mean enrichment was near 0, with high reproducibility between biological replicates (Fig S3B). Prior density centrifugation of BR-bodies comparing BR-body + and BR-body – strains found they were enriched in mRNAs, small non-coding RNAs, and antisense RNAs while depleted of tRNAs and rRNAs^16^. Our proximity labeled RNA samples showed mRNAs and small non-coding RNAs were both enriched, while antisense appeared to be enriched, however, the p-value was below a 95% confidence interval (p=0.1), while tRNAs and rRNA were not enriched (Fig 4C). Interestingly, while 16S and 23S rRNA were not enriched, 5S rRNA was enriched, in-line with RNase E’s known role of 5S rRNA processing^32,33^ and the colocalization of BR-bodies adjacent to rRNA loci in vivo^34^. Importantly though, while centrifugation-based isolation of BR-bodies required 200 mL of cell culture and 5 days of hands-on work to isolate the BR-body enriched RNAs^31^, APEX2 proximity labeling required only 4 mL of cell culture and 3 days of hands-on work, making it higher throughput.

**Figure 4.**
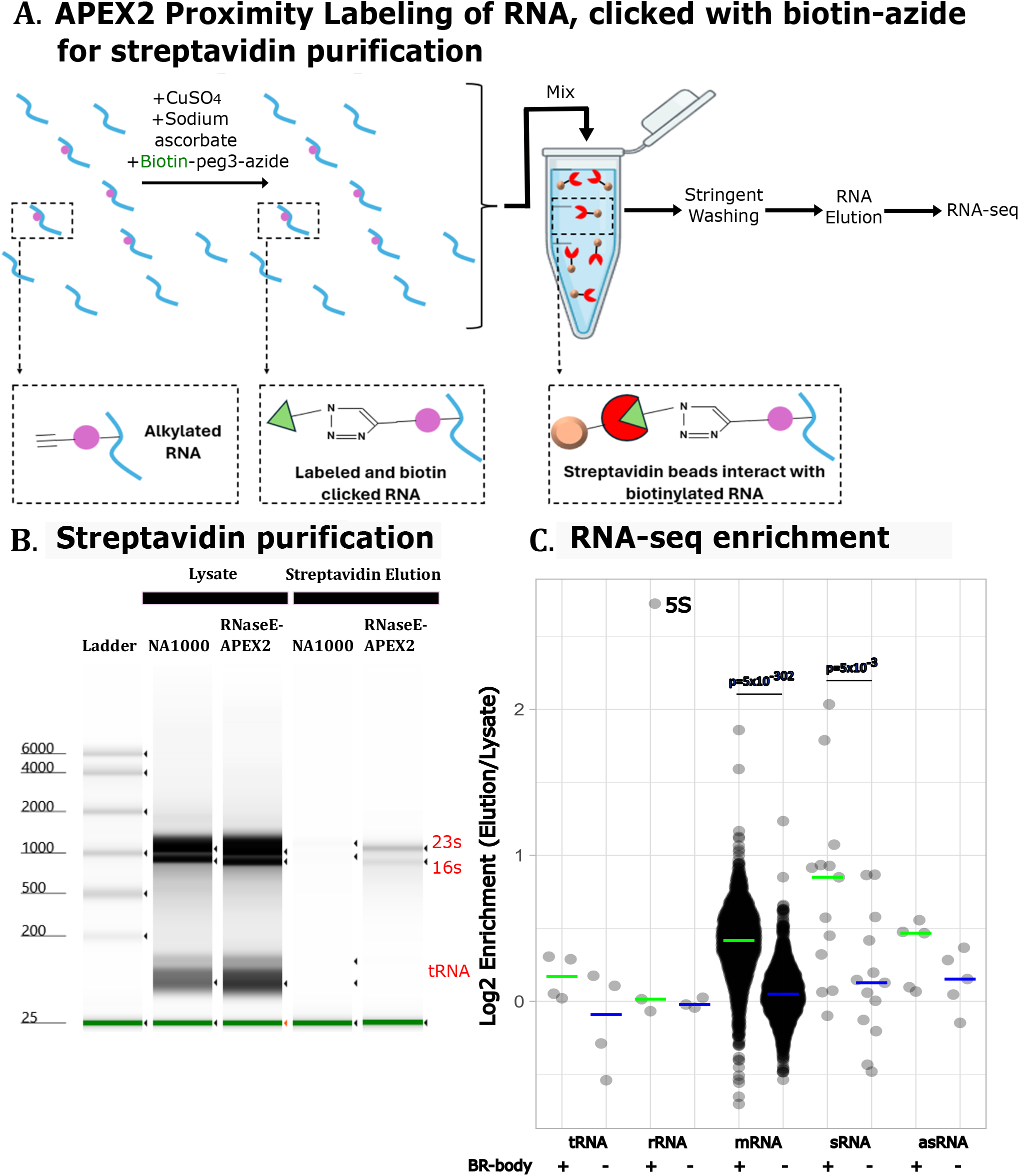
APEX2 proximity labeled RNA can be isolated by streptavidin purification. A) Schematic of the conjugation of biotin-azide to clicked alkyne-phenol and the resulting streptavidin purification. B) Streptavidin purification of biotinylated RNAs. Tapestation RNA profiles of the lysates and elution fractions of biotinylated proximity labeled RNAs. The bottom two bands are tRNAs, which are known to be highly depleted in BR-bodies from differential centrifugation-based isolation of BR-bodies^16,31^. C) RNA-seq enrichment analysis of proximity labeled RNA from BR-body + (RNaseE-APEX2, JS767) and BR-body – (RNaseEΔCTD-APEX2, JS801) cells. Log2 ratio of elution/lysate from each sample is plotted for each RNA quantified. Colored horizontal bars indicate the median values for each distribution. T-test with uneven variance was used to compare the distributions of each RNA type, and RNA types with p<0.05 are shown.

## Discussion

RNP complexes are important regulators of RNA biology, including transcription, RNA processing, transport, translation, and decay^4,35–37^. In addition to RNP complex formation, these complexes have been increasingly found to localize into biomolecular condensate structures, which help to facilitate the spatial organization of the stages in the mRNA life cycle via phase-separation^4,13,16,38–41^. As realizations that RNA localization has become increasingly important in bacteria, methods to identify the population of localized RNAs have been developed, yet these methods have been limited due to the long timescale of the procedures compared to the short mRNA lifetimes in bacteria. APEX2 proximity labeling can rapidly label RNA and can be easily genetically fused to genes whose proteins have known patterns of subcellular localization. APEX2 proximity labeling of RNA requires small amounts of cells, can be completed in a few hours by a single scientist, and yields sufficient RNA for downstream analysis. The reactivity radius of APEX2 is estimated to be 10-20nm^42^, suggesting it gives high spatial precision. Genetic fusion of APEX2 is easy to generate, APEX2 RNA labeling works across heme-producing bacteria (C. crescentus, B. subtilis, and E. coli), and APEX2 is similar in size to a fluorescent protein and has no observable functional impacts when fused to RNase E, so we anticipate that this is unlikely to disrupt target protein function. The flexibility of click chemistry allows the conjugation of a wide array of clickable fluorophores or clickable affinity substrates as biotin or azide-resin. Altogether, APEX2 proximity labeling of RNA will help to accelerate the discovery of localized RNAs in bacteria.

## Author contributions

HY performed RNA labeling experiments. JMS and HY designed the study and wrote the paper. HY generated C. crescentus APEX2 fusions strains. DK generated B. subtilis APEX2 fusions strains. ES, OG, and OAC generated E. coli APEX2 fusion strains.

## Acknowledgements

We thank Tamara Hendrickson for equipment, reagents, and hosting the Schrader lab after it was destroyed in a fire. We thank Dr. Anat Nussbaum-Shochat for technical help in the construction of ES413 E. coli strain. NIH grants R35GM124733 to JMS and R35GM131783 to DBK. WSU Career Chair Award to JMS. IU startup funds to JMS. NIH T32GM142519-03 to HY. Research in the OAC lab was supported by the Israel Science Foundation (ISF) founded by the Israel Academy of Sciences and Humanities (grant no. 1274/19). OAC is an incumbent of the Dr. Jacob Grunbaum Chair in Medical Sciences.

## Notes

### Competing Interest Statement

The authors have declared no competing interest.

### Summary of Updates

The prior version had listed an incorrect DNA sequence, so we have corrected it.

