## Supplementary figures and images for "APEX2 proximity labeling of RNA in bacteria"

### Fig S1

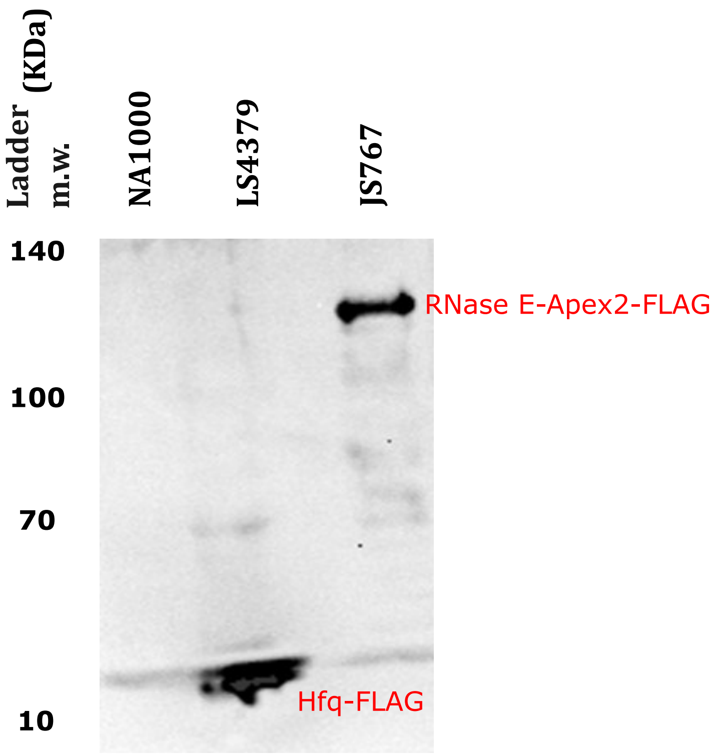

### Fig S2

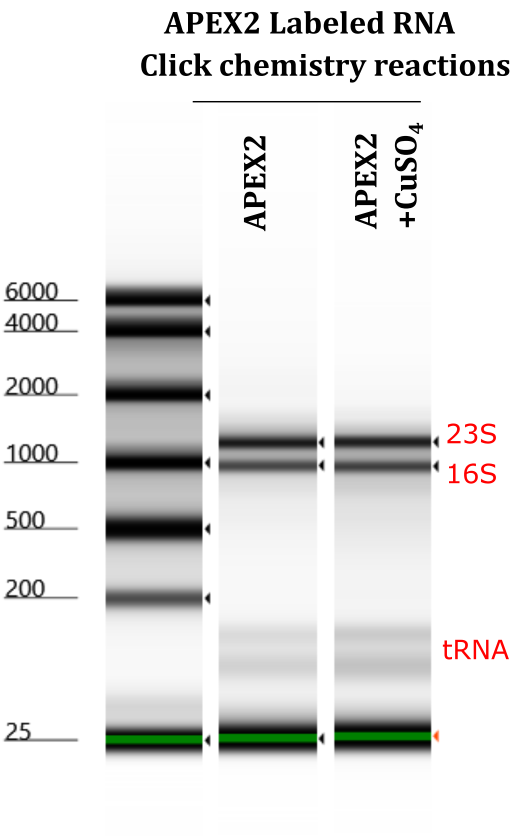

### Fig S3

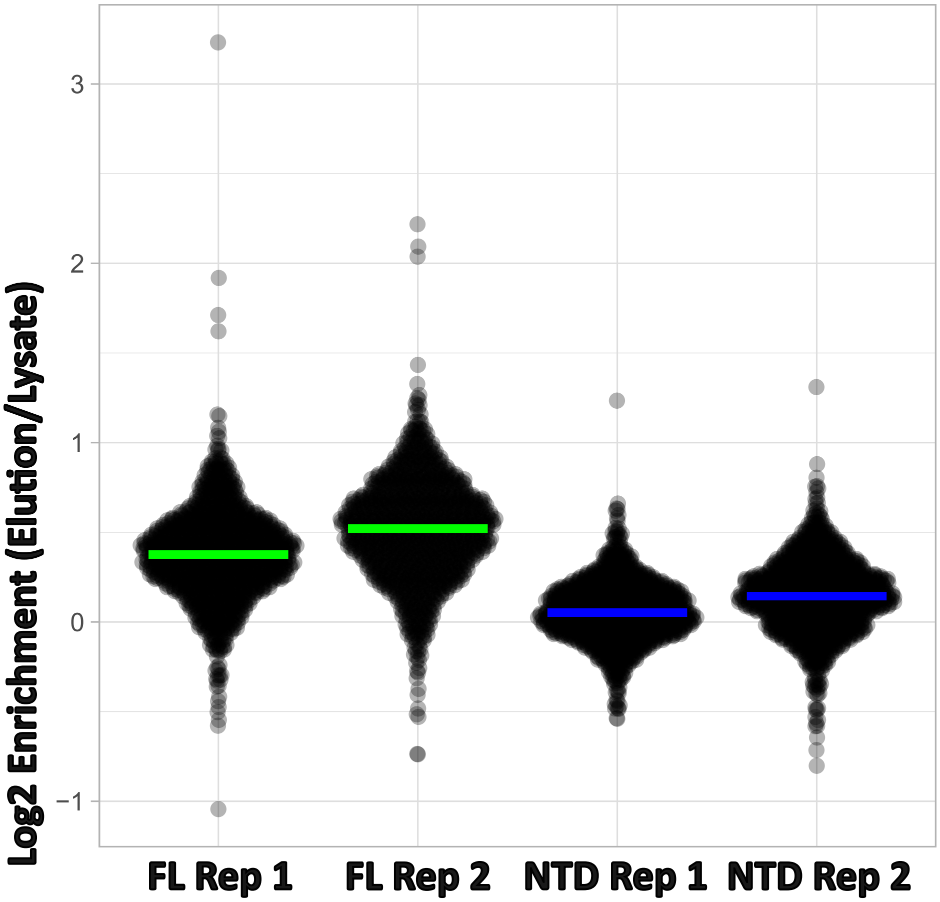

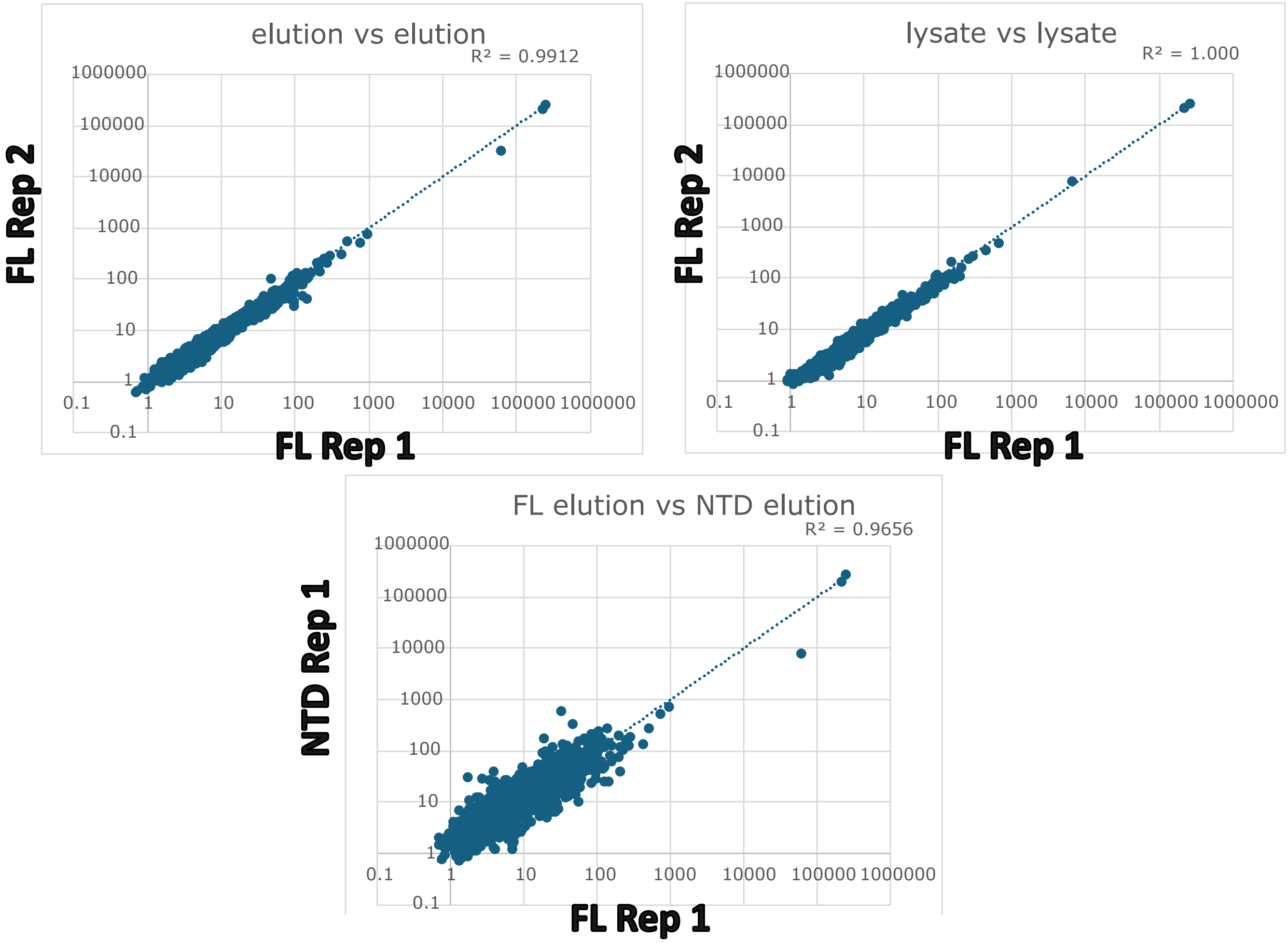
